# Epithelial innate immune sensing of pneumococci is inherently restricted to a small cellular minority across species and infection niches

**DOI:** 10.64898/2026.06.10.731054

**Authors:** Vikrant Minhas, Vincent de Bakker, Kin Ki Jim, Jun Kurushima, Joern Pezoldt, Monica Rengifo-Gonzalez, Camilla Ciolli Mattioli, Roi Avraham, Bart Deplancke, Jan-Willem Veening

## Abstract

*Streptococcus pneumoniae* colonises the nasopharynx asymptomatically yet causes life-threatening invasive disease. How it navigates early epithelial immune surveillance to cause disease remains unclear. Conventional innate immune models predict coordinated, population-wide epithelial responses to bacterial infection. Using single-cell RNA sequencing, RNA fluorescence in situ hybridization and in vivo mouse and zebrafish models, pneumococcal infection is instead shown to activate innate immune genes including chemokine, NF-κB regulatory, and prostaglandin pathway genes, in only 1-4% of lung epithelial cells. This restriction is seemingly pneumococcal-specific as *Escherichia coli* triggers responses in over 40% of the same cells. Strikingly, nasopharyngeal epithelial cells show complete immune silence to pneumococci while responding robustly to *E. coli* and *Staphylococcus aureus*, suggesting niche-specific immune evasion. Additionally, pharmacological inhibition of COX-2 significantly increased mortality in a zebrafish meningitis model, identifying prostaglandin signalling as a protective host response during invasive disease. Competence-associated surface remodelling contributes modestly and incrementally to immune restriction, while the predominant dampening is competence-independent. These findings challenge canonical epithelial immunity models against bacterial infection and provide a cellular framework for understanding pneumococcal commensalism and pathogenesis.

**Significance statement:** Classical innate immune models predict that bacterial infection triggers coordinated, population-wide transcriptional responses across the epithelium. Using single-cell RNA sequencing, RNA fluorescence in situ hybridization, and in vivo zebrafish and mouse models, we show that *Streptococcus pneumoniae*, responsible for over one million deaths annually, activates innate immune genes in only 1-4% of lung epithelial cells. *Escherichia coli* triggers responses in over 40% of the same cells, demonstrating this restriction is pneumococcal-specific. Nasopharyngeal epithelial cells, the bacterium’s primary colonization niche, show complete immune silence to pneumococci, suggesting niche-specific evolutionary adaptation. The prostaglandin pathway is identified as a protective host response during invasive disease. These findings challenge canonical models of epithelial immunity and provide a cellular framework for pneumococcal commensalism and pathogenesis.

## Introduction

*Streptococcus pneumoniae* (the pneumococcus) is a Gram-positive bacterium found in the upper respiratory tract where it lives as a commensal, yet it is also a leading cause of pneumonia, septicaemia, and meningitis, responsible for approximately one million deaths per year (1, 2). This paradoxical duality, asymptomatic nasopharyngeal carriage alongside the capacity for life-threatening invasive disease (3), raises a fundamental question about how the bacterium navigates host immunity during both colonization and infection. Conventional innate immune models predict that pathogen-associated molecular patterns (PAMPs) are detected by pattern recognition receptors expressed broadly across epithelial populations, triggering a coordinated, population-wide transcriptional response (4, 5). Whether pneumococcal infection conforms to this model, or whether epithelial immune activation is instead restricted to a minority of cells, has not been directly examined at single-cell resolution. Such restriction would be unexpected from a canonical danger-sensing perspective, yet would have profound implications for understanding how pneumococci persist asymptomatically in the nasopharynx and transition to invasive disease.

*S. pneumoniae* is naturally transformable and can assimilate exogenous DNA, in a phenomenon generally known as competence (6–8). Competence for natural transformation is a crucial mechanism of genome plasticity and is principally responsible for the acquisition and spread of antibiotic resistance as well as virulence factors such as the capsule (9, 10). Activation of competence upregulates a significant proportion of the pneumococcal genome with only a fraction of those genes required for DNA uptake and integration (11). In line with roles outside of transformation, competence has been shown to be activated during infection of human epithelial cells (12), invasive disease in mouse models (13, 14), and in a zebrafish infection model (15). The development of competence is a crucial pathogenic factor in pneumococcal meningitis (14, 15), and has been shown to remodel the cell wall exposing virulence factors by CbpD-based capsule shedding and a ComM-mediated block in division (Fig. 1A) (16, 17). Despite these advances, how competence influences host epithelial sensing at the level of individual cells remains unknown.

**Figure 1.**
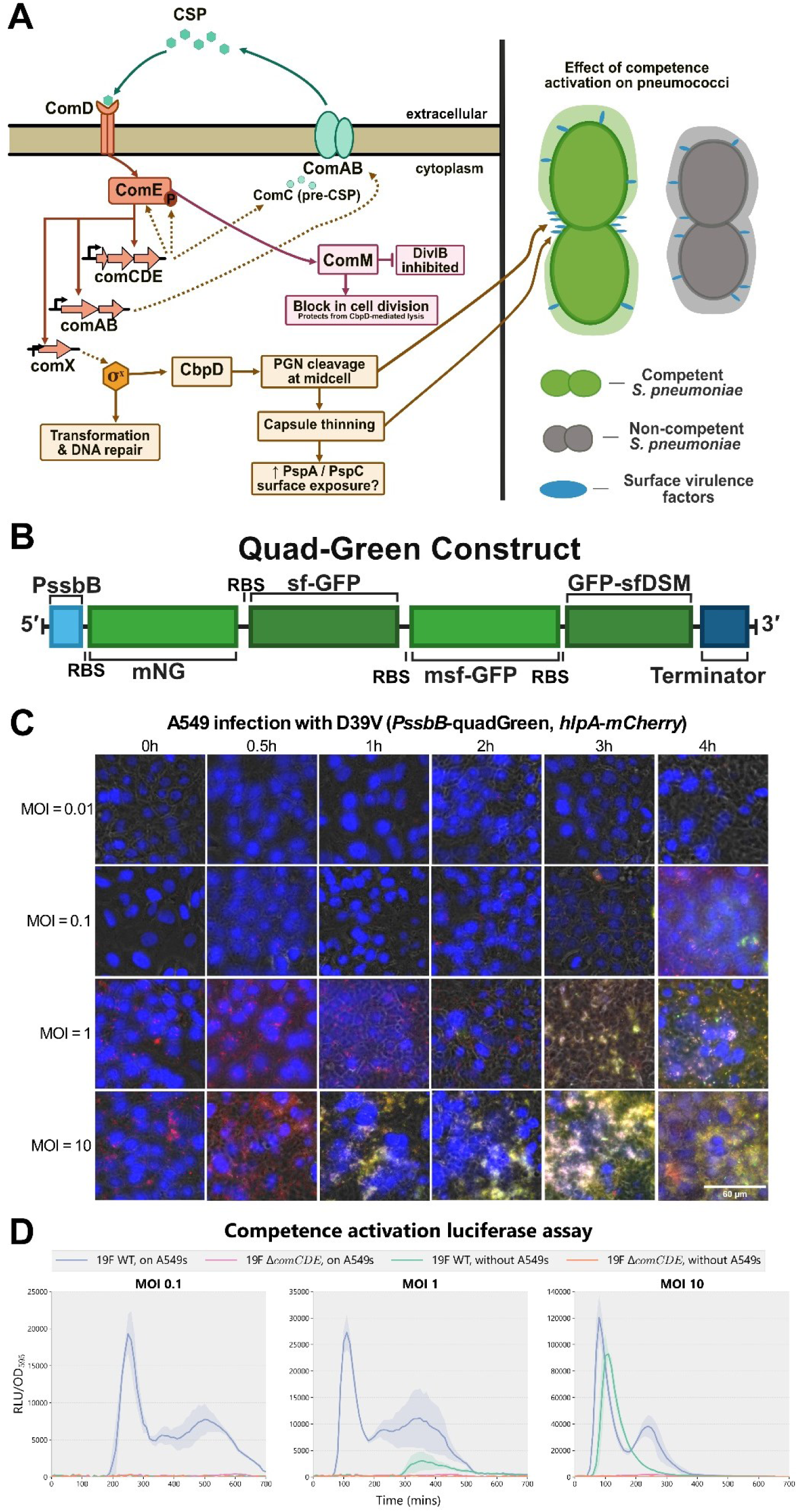
Pneumococci activate competence during lung epithelial infection in a heterogeneous manner. **A**) Schematic overview of the competence signalling pathway in *S. pneumoniae* and its downstream effects. Extracellular CSP is sensed by the ComD/ComE two-component system, leading to upregulation of *comCDE* and *comAB*, activation of ComX, and induction of late competence genes including DNA repair and transformation machinery. Competence activation also results in peptidoglycan (PGN) cleavage at midcell via CbpD, capsule thinning, increased surface exposure of PspA/PspC and inhibition of DivIB/a block in cell division. The right panel illustrates the morphological and surface differences between competent (green) and non-competent (grey) *S. pneumoniae*, highlighting increased exposure of surface virulence factors during competence. **B)** Schematic of the quadGreen construct, in which four green-fluorescent reporters that have been codon optimized for pneumococci (*mNG*, *sf-GFP*, *msf-GFP*, and *GFP-sfDSM*) are encoded in a single polycistronic operon downstream of the *PssbB* late competence promoter. **C)** A D39V pneumococcal strain containing a quadGreen fluorescence cassette coupled to the late competence promoter P*ssbB* and an *mCherry* fluorescence cassette attached to the constitutively expressed histone-like protein *hlpA* was inoculated with A549 cells at MOIs of 0.01, 0.1, 1, and 10. Nuclei of A549 cells were stained with DAPI (blue). Fluorescent images were taken at multiple timepoints during the 4h infection (see Materials and Methods for more details). **D**) A pneumococcal serotype 19F strain and an isogenic Δ*comCDE* mutant, both containing the *luc* gene attached to the late competence promoter P*ssbB*, were inoculated with or without A549 cells at MOIs of 0.1, 1, and 10. Luciferase activity (RLU/OD595) was measured over time to quantify competence induction. Three replicates for each condition and the standard error of the mean (SEM) are shown in the shaded area.

In general, many mechanisms that allow *S. pneumoniae* to cause invasive disease remain undetermined. Addressing this gap will require understanding the heterogeneous, single-cell level interactions between host and pathogen. While bulk RNA-seq analyses have shown that pneumococcal infection greatly changes the transcriptome of both pathogen and host (12, 18, 19), these approaches obscure heterogeneity within these interactions. Spatially or numerically restricted immune activation may be required to limit cellular damages following inflammation. Increasing evidence indicates that infection outcomes are strongly shaped by heterogeneity within both host and pathogen populations (20, 21). Single-cell transcriptomic approaches now enable direct analysis of heterogeneous host responses during infection, providing new insight into how individual cells contribute to tissue-level outcomes (22).

To directly address whether pneumococcal infection triggers a coordinated or heterogeneous epithelial response at single-cell resolution, we applied single-cell RNA sequencing (scRNA-seq) to human lung epithelial cells infected with GFP-expressing wild-type or competence-deficient pneumococci, enabling transcriptional profiling of infected and bystander cells separately. Contrary to canonical immune sensing models, we find that pneumococcal infection activates innate immunity in only a small minority of epithelial cells, a finding reproducible across independent experimental platforms and conserved across species, and that this restriction is most complete in the primary pneumococcal colonization niche.

## Results

### Pneumococci activate competence heterogeneously during lung epithelial infection

Previously, it has been shown by bulk dual RNA-seq that competence is activated during infection of A549 cells, the well characterized and widely used model for human type II epithelial cells, by the *S. pneumoniae* serotype 2 D39V strain (12). However, whether any heterogeneity occurs between host cells during competence activation was not examined. To obtain brightly fluorescent bacteria, visible with microscopy and flow cytometry, we constructed a so-called quadGreen cassette in which four different green-fluorescent reporters with alternative codon usages were cloned in a single polycistronic operon (Fig. 1B). To investigate potential heterogeneity, A549 cells were infected with a D39V pneumococcal strain, containing quadGreen and mCherry fluorescent cassettes, at multiple different MOIs, ranging from 0.01 to 10 for up to 4h (Fig. 1C). This quadGreen cassette was placed downstream of the P*ssbB* promoter, a late competence gene that is only activated upon competence (23). *mCherry* was fused to the histone-like protein *hlpA* under the control of a constitutive promoter. Hence, when competence is not induced, the bacteria are observed as red in fluorescence microscopy, but when competence is activated, bacteria also express green fluorescence. Indeed, it was observed, especially at higher MOIs, that many pneumococci had competence upregulated. However, bacteria with no competence upregulation can still be observed at all time points (Fig. 1C).

While D39V was selected for initial characterisation given its prior use in bulk dual RNA-seq studies of pneumococcal-epithelial infection (12), we used the serotype 19F strain to infect A549 cells, and for all subsequent experiments, given its clinical relevance as a globally dominant colonizer with high antimicrobial resistance rates (24–26). While the quadGreen cassette allows single-cell resolution visualization of competence heterogeneity by fluorescence microscopy, it is not suited for quantitative bulk assays. We therefore used a luciferase reporter under the same P*ssbB* promoter in the serotype 19F strain, which allows quantitative measurement of population-level competence induction kinetics. As was the case with strain D39V, contact with A549 cells increased luciferase levels in 19F and hence competence induction at all multiplicities of infection (MOI) tested (Fig. 1D, Supplementary Table 1). These data suggest that while pneumococci activate competence during infection of A549 lung epithelial cells, there is much heterogeneity in competence induction within these populations, similar to what was observed during infection in a zebrafish meningitis model (15).

### scRNA-seq shows only a subset of infected lung epithelial cells responding to pneumococcal infection

We reasoned that the transcriptional heterogeneity observed on the bacterial side may result in response variability on the host side too. To examine heterogeneous responses of host cells after infection with *S. pneumoniae*, A549 lung epithelial cells were infected with either wild-type serotype 19F pneumococci or an isogenic Δ*comCDE* knockout mutant, both constitutively expressing green fluorescence, at an MOI of 20. Since these epithelial cells were to be subjected to scRNA-seq following Fluorescence-Activated Cell Sorting (FACS), we used a relatively high MOI to ensure sufficient GFP-infected cells were sorted for this purpose. A completely naive uninfected control was also included. 2h after infection, infected and bystander (exposed to pneumococci but not directly infected at time of sorting) cells were sorted based on fluorescence (Supplementary Fig. 1) and these samples plus the naive control were subjected to 10x scRNA-seq (Supplementary Fig. 2A, Supplementary Tables 2-3). The 2h time-point was used since further time-points resulted in too many dead cells for efficient sorting.

We found that transcription profiles of cells across the samples (WT infected, WT bystander, Δ*comCDE* infected, Δ*comCDE* bystander and naive control) were generally homogeneous (Supplementary Fig. 2B-C), which is unsurprising given that all cells were derived from the same immortalized lung epithelial cell line. Of the three distinct clusters that we could identify, one was defined by above-average expression of NTS, which is associated with lung cancer (27), and another by upregulation of neutrophil-attracting CXCL cytokines including CXCL2, produced at inflammation sites (28) (Fig. 2A). Moreover, the latter cluster was characterized by a strong upregulation of general innate immune response genes, also including the cytokines CXCL1, CXCL3, CXCL5 and CXCL8 (29), the NF-κB and TNF regulators NFKBIA, NFKBIZ, IER3, TNFAIP3, ZFP36 and TNFAIP2 (30–33), as well as the immune regulators CEBPD and IRF1 (34, 35) (Fig. 2B). Further, naive cells, which were never exposed to bacteria, were almost entirely absent from this cluster (Fig. 2C). We therefore deduced that this cluster represents the cells that respond to the bacterial presence. The transcriptional response itself did not seem meaningfully different between challenged samples, implying that mere exposure to the bacteria was sufficient to trigger the response (Supplementary Fig. 2D). Although cells infected with competence-deficient bacteria mounted the immune response relatively more often (1.64%) than cells from the other samples (0.98% mutant bystander, 0.93% WT infected, 0.81% WT bystander), suggesting the mutants are more easily detected, these subpopulations were strikingly small across all samples (Fig. 2D, Supplementary Fig. 2E). This led to the interesting conclusion that only a fraction of human lung epithelial cells respond to pneumococcal infection after 2 hours, even at the relatively high MOI of 20. To validate the generality of our “small subpopulation immune response” observation, we revisited published scRNA-seq data of mouse lung tissue 12 hours post pneumococcal infection compared to a PBS-treated control (36). Despite the inevitably lower effective MOI, longer infection time, and general *in vivo* setting of that study, we indeed still found that specifically in the infected sample, only a subset of alveolar epithelial cells displayed elevated expression of several orthologs of these same innate immune response genes, most notably Cxcl2 and other chemokines (Fig. 2E), supporting the few percentage immune cell responding data from this study’s pneumococcal-lung epithelial scRNA-seq.

**Figure 2.**
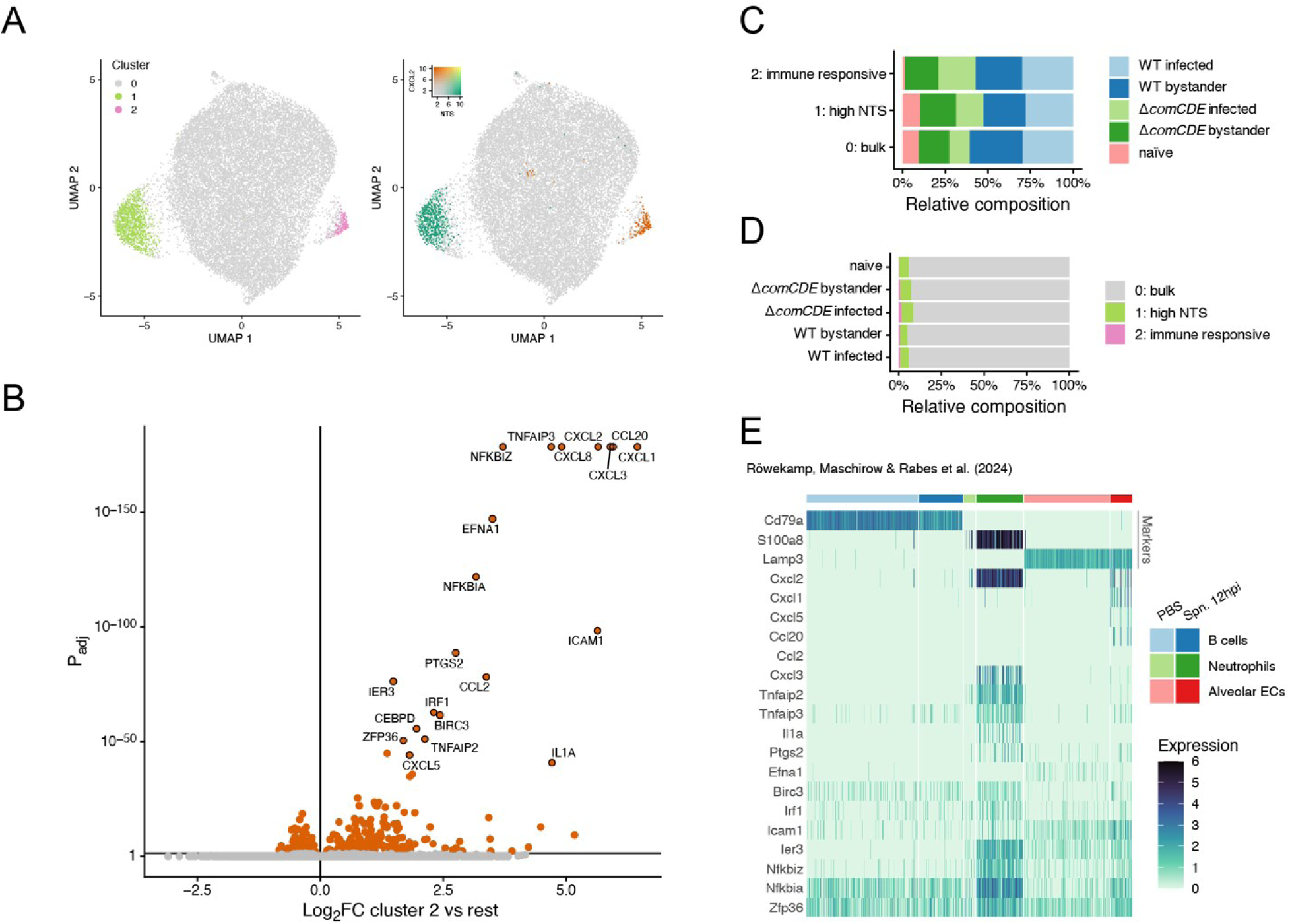
Single-cell RNA-seq reveals only a small subset of pneumococci-exposed lung epithelial cells mount an innate immune response two hours after infection. A) Cluster identities and normalized NTS and CXCL2 expression of cells in Uniform Manifold Approximation and Projection (UMAP) embedding. B) Differential expression between cluster 2, corresponding to immune-responsive cells, and others. C) Cluster composition in terms of samples that cells originate from. D) Fraction of cells assigned to each cluster per sample. E) Previously published scRNA-seq data show heterogeneity in the expression of various innate immune gene orthologs shown in Fig. 2B in mouse alveolar epithelial cells (in red) 12 hours post infection with *S. pneumoniae* compared to PBS treatment. Note also that the neutrophil population size increased considerably upon infection, as we have shown previously (18). The same global marker genes as in the original publication were used for cell type determination and are depicted in the top rows of the heatmap, as indicated. Columns represent individual cells in the respective cell type clusters. Expression was normalized with the variance-stabilizing normalization of the SCTransform() functionality in the Seurat R package. Data were obtained from Röwekamp, Maschirow & Rabes et al (36), NCBI GEO accession GSE236344.

### RNA FISH shows a subset of infected host cells activate cytokine genes during infection

To further validate the scRNA-seq results, which showed only a subset of A549 cells responding to pneumococcal infection, single molecule fluorescence in situ hybridization (smFISH) was performed. Here, A549 lung epithelial cells were infected with WT and isogenic Δ*comCDE* serotype 19F *S. pneumoniae* strains, under the same conditions as in the scRNA-seq experiments. These samples were then processed for smFISH targeting the constitutively expressed control gene GAPDH and innate immune genes CXCL2 and NFKBIA (Fig. 3A-C, Supplementary Table 4). In line with the scRNA-seq data, only a small fraction of cells activated innate immunity following pneumococcal infection. Specifically, 2.49% and 4.14% of A549 cells upregulated CXCL2, while 2.84% and 4.76% of A549 cells upregulated NFKBIA, 2 hours after infection with 19F WT and Δ*comCDE*, respectively. In contrast, uninfected A549 cells showed no expression of these cytokines (Fig. 3B). These values are consistent with the scRNA-seq data and further suggest that restriction of innate immune activation to a small epithelial minority is a robust, reproducible feature of the host response to pneumococcal infection, validated here by an entirely independent single cell analysis approach. As cells infected with competence deficient bacteria were more likely to activate the CXCL2 and NFKBIA genes compared to cells infected with WT infection, we tested if inducing competence externally through the addition of CSP-1 in WT pneumococci decreased the A549 CXCL2 response. Indeed, when WT pneumococci had competence induced externally, the percentage of cells expressing CXCL2 and NFKBIA dropped slightly further, to 1.80% and 1.87%, respectively, compared to when cell were infected with uninduced WT pneumococci that would only activate competence at later time points (Fig. 3B). To test whether heterogenous expression of the innate immune system in A549 cells is specific to pneumococcal infection, we also infected cells with *Staphylococcus aureus*. Interestingly, infection of A549s with *S. aureus* resulted in a higher percentage of A549s upregulating CXCL2 and NFKBIA, 7.14% and 7.51%, respectively, compared to *S. pneumoniae* infection at the same MOI (Fig. 3B), though the exact percentage of infected cells between the two strains was not determined. To assess if A549 cells are in principle all capable of activating the innate immune response, we infected A549 cells at a high MOI of *Escherichia coli,* which is known to trigger the innate immune response through its LPS (37, 38). Indeed, significantly more A549 cells infected with *E. coli* expressed CXCL2 and NFKBIA, 41.09% and 36.98%, respectively (Fig. 3B), showing that in principle many lung epithelial cells are capable of expressing these cytokines.

**Figure 3.**
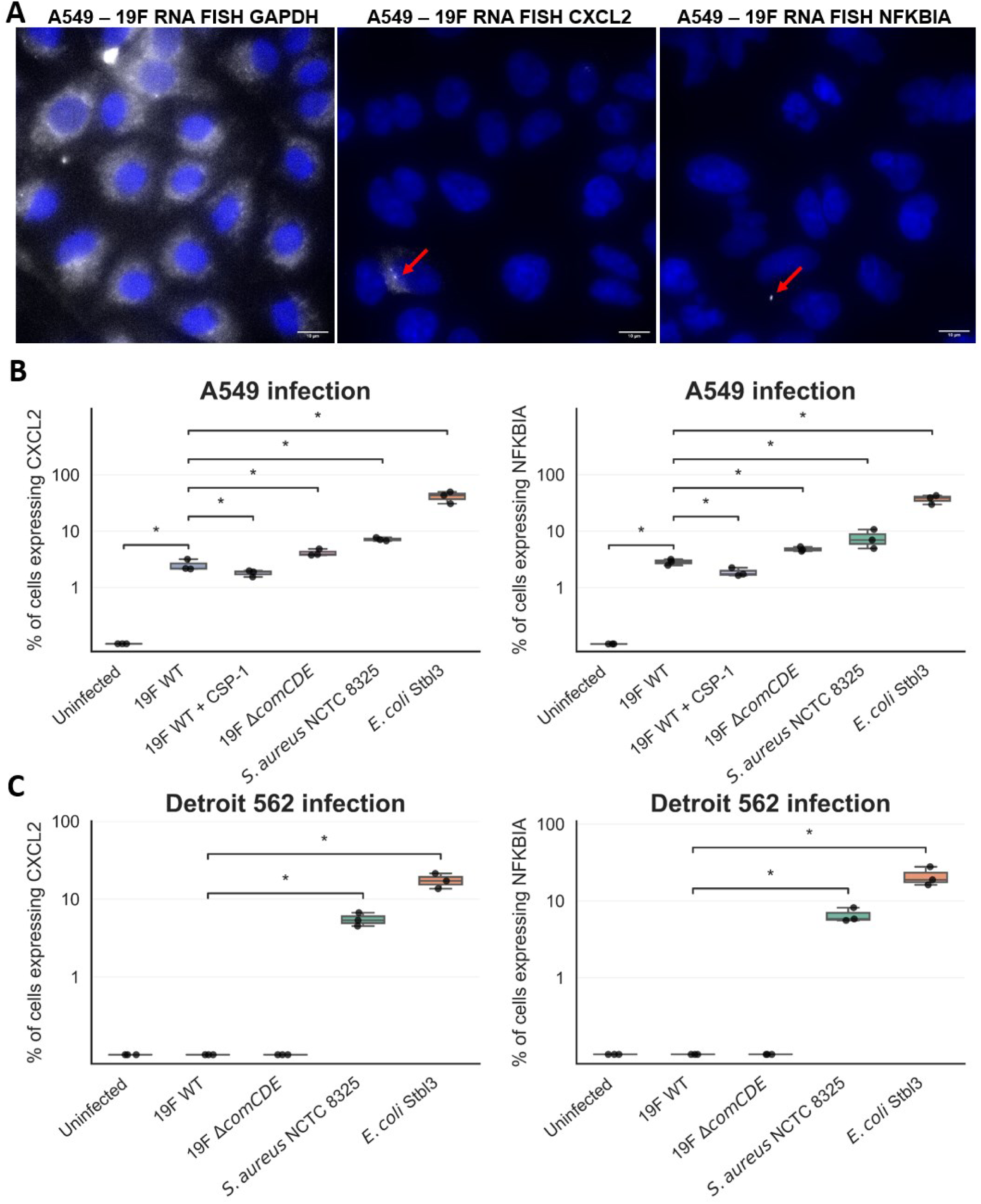
RNA FISH to examine CXCL2 and NFKBIA expression. A) Representative RNA FISH images of *S. pneumoniae* 19F infected A549 cells 2h post infection at an MOI of 20. Samples were stained with DAPI to visualize the nuclei in blue. The white dots represent the presence of either GAPDH, which acted as positive control, or CXCL2 and NFKBIA mRNA (indicated by the red arrow). B-C) To quantify the RNA FISH data, the percentage of host cells expressing CXCL2 was calculated from several images in triplicate experiments for A549 (B) or Detroit 562 (C) cells infected with the pneumococcal 19F WT (+ CSP-1 only for the A549 condition) and Δ*comCDE* strains as well as *S. aureus* NCTC 8325 and *E. coli* STBL3 strains. Data presented are the medians and interquartile range (*, P < 0.05, Mann-Whitney U test).

While human A549 are lung alveolar epithelial cells, the primary pneumococcal niche is the nasopharynx. To test whether bacteria also trigger an immune response in nasopharyngeal epithelial cells, *S. pneumoniae* 19F bacteria, *S. aureus* and *E. coli* were used to infect nasopharyngeal epithelial Detroit 562 cells. Interestingly, Detroit 562 cells did not activate CXCL2 or NFKBIA when infected with either strain of pneumococci, while infection with *S. aureus* and *E. coli* showed 5.50% and 17.34% of cells expressing CXCL2, respectively, and 6.50% and 20.89% of cells expressing NFKBIA, respectively, 2h post-infection (Fig. 3C). This suggests that pneumococci are either not detected by nasopharyngeal cells or nasopharyngeal cells do not activate these genes when seeing *S. pneumoniae*. Alternatively, it could be envisaged that *S. pneumoniae* is capable of repressing CXCL2 and NFKBIA expression in nasopharyngeal cells during early colonization.

### Zebrafish embryo meningitis model validates innate immune activation and identifies a protective role of the prostaglandin pathway

To validate whether host responses identified in epithelial cells were conserved *in vivo*, we used the zebrafish embryo meningitis model to assess innate immune activation during invasive pneumococcal disease. The single-cell RNA sequencing above revealed that only a subset of infected epithelial cells mounted a pronounced inflammatory response, characterised by induction of innate immune signalling pathways (Fig. 2B). To test whether this immune response is also conserved in a zebrafish infection model, quantitative RT-qPCR was performed on whole zebrafish embryos on a selection of top differentially expressed genes from this inflammatory subset. Specifically, as CXCL2 itself lacks a direct zebrafish ortholog, *cxcl8a* was used as the functional equivalent for neutrophil chemoattraction. The genes *nfkbiaa*, *nfkbiz*, and *tnfaip3* were selected as conserved NF-κB pathway regulators identified in the scRNA-seq data, while *ptgs2a* and *ptgs2b* were prioritised given independent evidence for COX-2 induction during pneumococcal infection in human tissue (39, 40) and the functional relevance of this pathway tested below. Here, we confirmed induction of these genes (Fig. 4A, Supplementary Table 5), supporting that this response is preserved *in vivo* within the context of invasive pneumococcal disease. Notably, *cxcl8a*, encoding a key neutrophil chemoattractant, and *nfkbiaa*, a negative regulator of NF-κB signalling, were consistently more highly induced in Δ*comCD*E-infected embryos at both time points. Additionally, real-time fluorescence time-lapse imaging using Tg(mpx:GFP) zebrafish embryos with fluorescently labelled neutrophils showed clear neutrophil migration upon infection with either strain without a significant difference between them (Fig. 4B, Supplementary Table 6). Notably, both *ptgs2a* and *ptgs2b* were induced at 8 hpi across both strains, suggesting that PTGS2-dependent prostaglandin signalling is a key component of the host response during pneumococcal infection in the zebrafish model. COX-2 has previously been shown to be induced in lung epithelial cells and human lung tissue during pneumococcal infection, where it contributes to prostaglandin-mediated inflammatory regulation (39, 40). To directly test the functional relevance of this pathway, pharmacological inhibition of COX-2 activity using celecoxib resulted in increased mortality of infected zebrafish embryos (Fig. 4C), suggesting that PTGS2-dependent signalling plays a protective role during pneumococcal meningitis. Together, these findings do not find a strong role for pneumococcal competence in modulating early host-pathogen interactions but did identify a protective role of the prostaglandin pathway.

**Figure 4.**
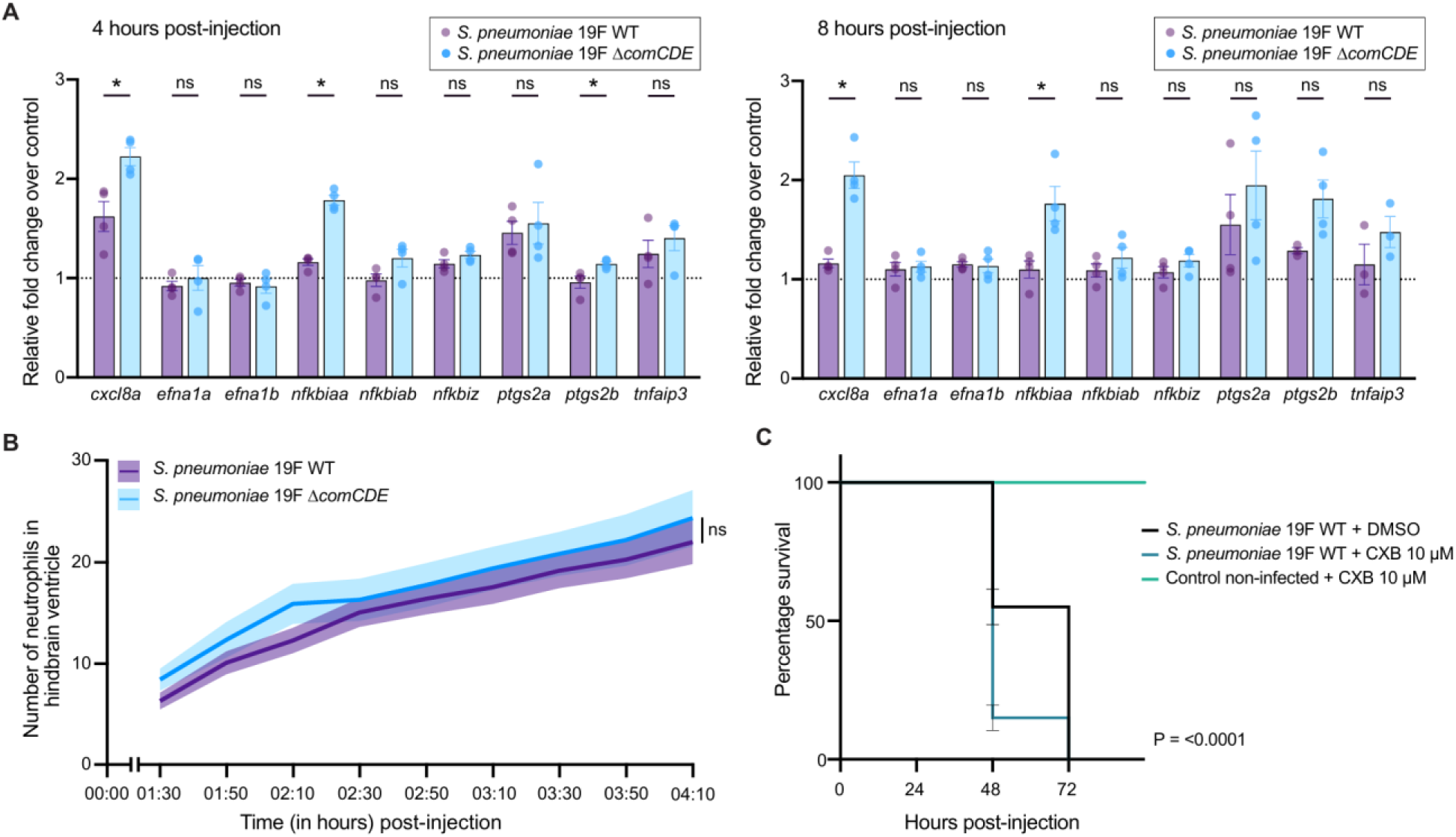
Validation of innate immune activation and identification of a protective role of the prostaglandin pathway in a zebrafish meningitis model. (A) Gene expression analysis of inflammatory markers during early stages of pneumococcal meningitis. RT-qPCR was performed on pools of 20 zebrafish embryos injected at 2 days post-fertilisation into the hindbrain ventricle with approximately 2,000 CFU of *S. pneumoniae* serotype 19F wild type (WT) or 19F Δ*comCDE*, or mock-injected controls, and collected at 4 and 8 hours post-injection (hpi). Relative fold change was calculated over non-infected controls. Data represent 3-4 biological replicates. Statistical comparisons between strains at each time point were performed using the Mann-Whitney U test; * = P < 0.05. (B) Neutrophil recruitment to the hindbrain ventricle was monitored over time by live fluorescence imaging in *Tg(mpx:*GFP) zebrafish embryos infected at 3 dpf with approximately 1,000 CFU of 19F WT or 19F Δ*comCDE* (n = 35 and 29, respectively, across two individual experiments). Imaging was initiated at 1.5 hpi and performed at 20-minute intervals. Notably, neutrophils are absent from the hindbrain ventricle in uninfected embryos, meaning that all neutrophils observed at this site are recruited in response to infection. Statistical analysis was performed using two-way repeated-measures ANOVA with Geisser-Greenhouse correction. Overall neutrophil numbers did not differ significantly between strains (F(1, 62) = 1.649, p = 0.2039), and no significant strain x time interaction was observed (F(8, 496) = 0.3250, p = 0.9565). Results are presented as mean ± SEM. (C) Survival analysis of 2 days post-fertilisation zebrafish embryos injected with approximately 400 CFU of *S. pneumoniae* 19F WT or vehicle control injected (control non-infected) into the hindbrain ventricle and treated with the COX-2 inhibitor celecoxib (CXB) at 10 µM. Data represent three biological replicates with 20 embryos per group (60 embryos in total per group) and were analysed using the log-rank Mantel-Cox test; *S. pneumoniae* 19F WT vehicle control vs *S. pneumoniae* 19F WT + 10 µM CXB, P < 0.0001.

## Discussion

Classical innate immune theory holds that epithelial cells, as a first line of defence, detect microbial invaders broadly through constitutively expressed pattern recognition receptors, mounting a coordinated transcriptional alarm across the exposed cell population (4, 5). Our data directly challenge this expectation for pneumococcal infection, that rather than a population-wide response, innate immune activation is confined to roughly 1-4% of lung epithelial cells within the first two hours of exposure (Fig. 2). This restriction is not due to limited epithelial immune capacity, as *S. aureus* and *E. coli* infection under identical conditions induced CXCL2 and NFKBIA expression in substantially larger fractions of cells (Fig. 3). Indeed, *S. pneumoniae* occupies a paradoxical position, living as a common commensal in the nasopharynx while remaining a leading cause of invasive disease (1), and this restriction of early epithelial signalling likely contributes to that duality. This cellular dichotomy, innate immune activation confined to a small epithelial minority during pneumococcal infection, contrasted with the broad population-wide response elicited by *E. coli*, is summarised schematically in Fig. 5. While bulk RNA-seq has previously shown how infection rewires host and bacterial transcriptomes (12, 18, 19), these population-level averages tend to mask interesting cell-to-cell variations. By using a single-cell approach, we show that the innate immune response in the epithelium is indeed heterogeneous. Importantly, epithelial immune heterogeneity was also shown in vivo in a murine model of pneumococcal pneumonia. Reanalysis of publicly available mouse lung scRNA-seq data revealed that pneumococcal infection induces innate immune gene expression in only a subset of alveolar epithelial cells (Fig. 2E), mirroring our in vitro observations (36). The concordance between a human epithelial cell line infected in vitro at high MOI and mouse alveolar epithelial cells infected in vivo with considerably lower bacterial burden argues strongly that this restricted response reflects a genuine biological feature of the host-pneumococcal interaction, rather than a consequence of experimental conditions, cell line artefact, or limited scRNA-seq sensitivity. Moreover, the 2-hour infection timepoint was selected to capture early transcriptional responses prior to substantial cell death, which precluded efficient sorting at later timepoints. This timepoint is consistent with the kinetics of early NF-κB dependent innate immune gene induction reported in bulk epithelial infection studies, and the concordance between our 2-hour in vitro data and the 12-hour in vivo mouse scRNA-seq dataset argues against the minority response being an artefact of the early time point studied. While the zebrafish data were generated at the whole-embryo level using bulk RT-qPCR and therefore cannot examine cellular heterogeneity, they provide evidence to suggest that the specific signalling pathways activated within the minority-responding epithelial population are biologically relevant during genuine invasive disease in vivo. Pharmacological inhibition of COX-2, encoded by a gene also induced in this minority cluster, significantly increased embryo mortality, further reinforcing that even the signalling output of a small responding minority can have measurable consequences for infection outcome.

**Figure 5.**
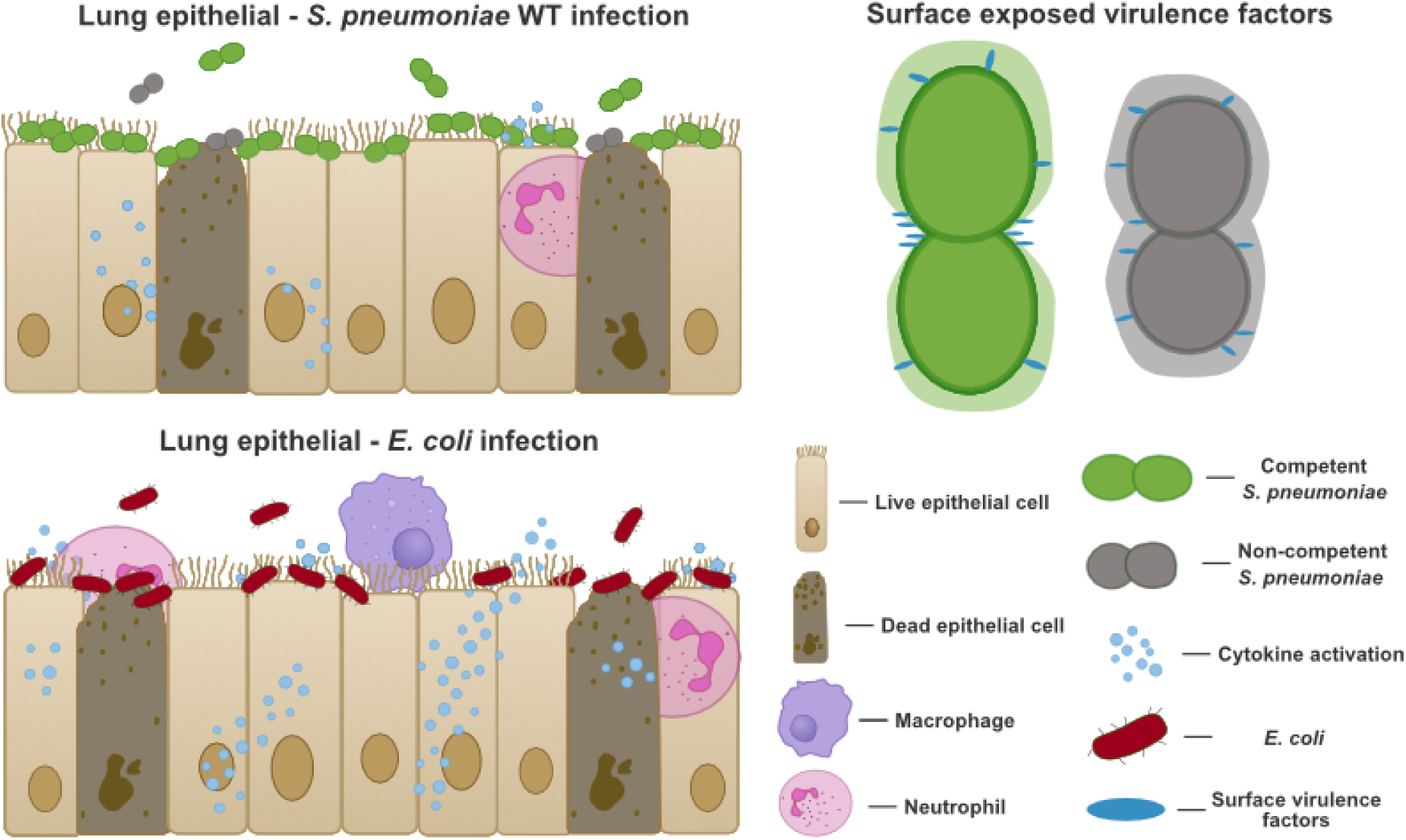
Model of epithelial innate immune heterogeneity during pneumococcal and *E. coli* infection. Top: During *S. pneumoniae* WT infection of lung epithelial cells, innate immune activation, represented by cytokine induction, is restricted to a small minority of cells (∼1-4%), with the majority of the epithelial monolayer remaining transcriptionally quiescent. Immune cell recruitment is correspondingly limited. Competent (green) and non-competent (grey) pneumococci both associate with the epithelial surface, with competent bacteria exposing additional surface virulence factors that may contribute incrementally to immune dampening. Bottom: In contrast, infection with *E. coli* elicits widespread cytokine activation across the epithelial population, with robust immune cell recruitment including neutrophils and macrophages and substantially greater epithelial cell death. Together, these panels illustrate that the restriction of epithelial innate immune signalling to a small cellular minority is a feature specific to pneumococcal infection, rather than a general property of epithelial-bacterial encounters.

Perhaps the most striking finding of pneumococcal-specific immune dampening in this study is the complete absence of CXCL2 and NFKBIA induction in nasopharyngeal epithelial cells, the primary colonization niche of the pneumococcus. Cells that mount robust responses to *S. aureus* and *E. coli* fail entirely to activate these innate immune genes in response to *S. pneumoniae*, irrespective of whether bacteria are competence-proficient or competence-deficient. This suggests that immune quiescence in the nasopharynx is not a property of the host cells, which are clearly capable of responding, but a consequence of pneumococcal-specific evasion mechanisms operating in that niche independently of competence. Notably, *S. aureus*, which colonizes the anterior nares rather than the nasopharynx (41), does trigger a cytokine response in Detroit 562 nasopharyngeal cells. This niche specificity, immune silence for a dedicated nasopharyngeal colonizer, immune activation for a bacterium of an adjacent but distinct mucosal site, suggests that the quiescence observed here reflects specific evolutionary adaptation by *S. pneumoniae* to its colonization niche, rather than a general property of commensal bacteria (1, 38, 39). Indeed, several previously characterised pneumococcal mechanisms contribute to the generally muted epithelial response observed in this and other studies. The major virulence factor pneumolysin induces host epigenetic modifications that broadly dampen innate immune gene expression (42), an effect that may involve COMMD2-mediated degradation of the NF-κB subunit p65 (43). Phosphorylcholine moieties on the pneumococcal cell surface mimic host molecules and suppress lectin pathway activation (44), while choline-binding proteins including CbpA directly interfere with complement recognition and neutrophil recruitment (45, 46). These mechanisms collectively provide potential context for why pneumococcal infection is globally less immunostimulatory than infection with *E. coli* or *S. aureus*. It will be interesting to determine whether there is a correlation between globally dominant serotypes and their immune-modulating capacities, as has been demonstrated for disease tropism and serotype-dependent differences in macrophage and inflammasome activation (47–49).

However, explaining why infection globally produces a quieter response is distinct from explaining why, within the same infection, 96-99% of cells are not expressing an innate immune response, while 1-4% mount a robust response. The present data do not resolve this second, more specific question. Several non-mutually exclusive mechanisms could account for this within-population restriction. Firstly, epithelial cells are known to be heterogeneous in their expression of pattern recognition receptors including Toll-like receptors and NOD-like receptors, such that individual cells may vary substantially in their intrinsic detection capacity (4, 5). Secondly, stochastic dynamics of NF-κB signalling, well documented at single-cell resolution, means that even nominally uniform stimulation can produce binary, all-or-nothing transcriptional responses in individual cells, with only those crossing an activation threshold producing detectable output (50, 51). Thirdly, actual per-cell bacterial encounter may vary substantially even at high nominal MOIs, such that many cells never achieve sufficient direct bacterial contact to trigger signalling. This is supported by our own sorting data, which show that even at MOI 20, only a minority of cells carried detectable GFP-positive bacteria at the time of sorting, the same observation that necessitated using MOI 20 in the first place. The effective per-cell encounter rate, rather than the population-level MOI, may therefore be the relevant parameter, consistent with the observed heterogeneity in competence induction shown in Fig. 1 where bacteria at the same population density activate competence at variable rates. Fourthly, the pneumococcal immune evasion mechanisms described above may lower the probability of detection for most cells while being insufficient to prevent activation in the rare minority of host cells that encounter a particularly high local bacterial load or are intrinsically more sensitive. The present data cannot distinguish between these possibilities. Doing so will require experimental designs that directly quantify per-cell bacterial burden during infection and correlate it with single-cell transcriptional output, or that characterise pattern recognition receptor expression heterogeneity across the epithelial population. Consistent with this, both bystander and infected cells showed immune activation (Fig. 2D), suggesting that physical bacterial contact at the moment of sorting is not the sole determinant of transcriptional response as epithelial cells may have encountered bacteria or bacterial products earlier during infection, with responses persisting after physical dissociation (20, 22).

Our data also suggest that pneumococcal competence may act as a potential modulator of epithelial immune activation. Although global host transcriptional profiles were broadly similar following infection with wild-type or Δ*comCDE* pneumococci, the fraction of immune-responsive epithelial cells was consistently, though only slightly, higher upon infection with the competence-deficient mutant. Deletion of the *comCDE* operon resulted in a modest increase in the fraction of immune responding cells, observed both by scRNA-seq and RNA FISH (Fig. 2D, Fig. 3B); however, these differences were limited in magnitude with only small numbers of cells making up these populations. Given that only a small subset of epithelial cells respond at this early stage, even modest shifts in responder frequency may potentially have substantial downstream consequences (20). How the inability to activate competence can influence epithelial immune signalling remains unclear. Competence-associated remodelling of the pneumococcal cell envelope provides a plausible mechanistic basis for immune dampening. Competence remodels the cell wall and increases surface exposure of key virulence factors (16, 52). Several of these factors interfere with complement activation and innate immune recognition, including pneumococcal surface protein A and choline-binding protein A (45, 53). In addition, modifications to pneumococcal phosphorylcholine are known to influence host detection via the lectin pathway (44, 54). Speculatively, competence-associated surface remodelling may therefore make a modest, incremental contribution to attenuating early epithelial innate immune recognition, consistent with increased surface exposure of known immune-modulatory virulence factors including PspA and CbpA (45, 53). However, the predominant immune dampening, particularly the complete immune silence in nasopharyngeal cells and the broadly restricted epithelial response in lung cells, appears to be competence-independent and likely reflects the broader pneumococcal-specific mechanisms described above. What drives the primary restriction of epithelial immune signalling to a small cellular minority remains the central open question this study raises.

Beyond the specific context of pneumococcal infection, these findings contribute to an emerging picture in infection biology in which tissue-level immune outcomes are not determined by uniform population responses but by the behaviour of small, responding cell minorities embedded within largely silent populations. Related heterogeneity has been described in macrophage responses to Salmonella, where variability in bacterial PhoPQ activity across the infecting population drives bimodal Type I interferon induction in individual host cells (20). Though in that case pathogen-side heterogeneity is the primary and sufficient driver, whereas in the present study the restriction appears likely to reflect pneumococcal-specific evasion mechanisms, but the specific bacterial factors responsible for the predominant, competence-independent dampening remain to be identified. Moreover, stochastic, all-or-nothing NF-κB dynamics documented in non-infectious inflammatory contexts may act in concert with bacterial evasion, shaping which of the rare host cells receiving sufficient signal ultimately cross the activation threshold to produce a detectable transcriptional response (50, 51). The present data extend this principle to epithelial innate immunity during bacterial colonization of a mucosal surface, a context where the stakes of the response threshold are particularly high, since excessive signalling risks inflammatory pathology while insufficient signalling permits pathogen expansion. Whether the 1-4% responder frequency identified here represents a conserved quantitative feature of mucosal epithelial immunity, or whether it varies with pathogen, niche, and host state, is to be determined, with implications well beyond pneumococcal biology.

## Methods

### Bacterial and epithelial cells culture conditions

All bacterial strains used in this work are listed in Supplementary table 1. *S. pneumoniae* strains were grown from a frozen culture in a liquid semi-defined casein-based C+Y medium (pH 6.8), supplemented with yeast extract (Sigma-Aldrich), *S. aureus* strains were grown in Tryptic Soy Broth and *E.coli* strains were grown in Lysogeny Broth at 37°C (52). Strains were grown from a starting optical density (OD_600_) of 0.01 until the appropriate OD for subsequent experiments. The human type II lung epithelial cell line, A549 (ATCC® CCL-185) and human nasopharyngeal cell line Detroit 562 (ATCC® CCL-138) were routinely cultured in DMEM (Dulbecco’s modified Eagle medium-nutrient mixture) F-12 with GlutaMAX (Life Technologies) supplemented with 10% (v/v) fetal bovine serum (FBS; VWR), 25mM HEPES buffer (Sigma-Aldrich) and Penicillin (50units/ml) - Streptomycin (50μl/ml) (Gibco). The cell culture was maintained at humidified 5% (v/v) CO_2_ atmosphere at 37°C on sterile 75cm^2^ cell-culture treated flasks (Corning), with Trypsin-EDTA solution (Sigma-Aldrich) used for cell detachment and a sub-culturing dilution of 1:25.

All primers used in this work are listed in Supplementary table 2. The quadruple-green construct was cloned by Golden Gate Assembly using AarI. The plasmid pVL2132, which carries de mNeonGreen-Opt sequence, was used as a backbone and was amplified with oligos OVL1584 and OVL1585. The sf-GFP-Sp-sequence was amplified from pVL306 (pJWV102) using oligos OVL1582 and OVL1583. The msf-GFP-OPT sequence was amplified from pVL2130 using oligos OVL1588 and OVL1589. The msf-GFP-DSM sequence was amplified from VL2129 using oligos OVL1586 and OVL1587. The resulting Golden Gate Assembly product was transformed in *E. coli* Stbl3 before being inserted into the *Plac* region of *S. pneumoniae* D39V and 19F strains. The *ssbb-luc-kan* cassette was transferred to the pneumococcal 19F strains from a D39V strain that was constructed by Slager et al (55), while the *comCDE*::*ery* 19F strains were constructed by inserting an erythromycin resistance cassette in place of the *comCDE* operon via homologous recombination directly into the pneumococci.

### Competence assays

For the luciferase activity assay, 200μl of A549 cells in DMEM culture medium at a density of 1.5x10^5^ cells/ml were seeded onto cell-culture treated 96 well trays (Corning Costar). The day after seeding, A549 cells in each well were rinsed with 1x PBS (pH 7.4) and 200μl of RPMI 1640 without phenol red (Life Technologies) supplemented with 1% (v/v) FBS was added. The following day, *S. pneumoniae* strains VL3894 and VL3889, which contain a transcriptional fusion of the firefly luciferase gene (*luc*) with the late competence gene *ssbB,* were grown in C+Y medium pH 7.8 +/- 0.05 (permissive conditions for natural competence induction), and upon reaching OD_600_ ∼0.1 (exponential phase), bacteria were spun down and resuspended to OD_600_ ∼0.01 in C+Y medium + 0.01mg/ml luciferin. 200μl of samples were added in triplicate to wells of the 96 well tray with or without A549s. The *S. pneumoniae* strains were cultured in a Tecan Infinite F200 PRO allowing for real-time monitoring of competence induction *in vitro*. Growth (OD_595 nm_) and luciferase activity (RLU) were measured every 10 min for 12 h. Expression of the *luc* gene (only if competence is activated) results in the production of luciferase and thereby in the emission of light. Three replicates for each condition and the standard error of the mean (SEM) are shown.

For the fluorescence microscopy assay, a 12mm coverslip (Menzel^TM^) was added to each well of a 24 well trays (Corning Costar) then 1 ml of A549 cells in DMEM culture medium at a density of 2x10^5^ cells/ml were seeded onto each well. The day after seeding, A549 cells in each well were rinsed with 1x PBS (pH 7.4) and 1ml of RPMI 1640 without phenol red (Gibco™) supplemented with 1% (v/v) FBS was added. The following day, *S. pneumoniae* strains were grown in C+Y medium, and upon reaching OD_600_ ∼0.2 (exponential phase), bacteria were spun down and resuspended in infection medium, RPMI 1640 without phenol red supplemented with 1% (v/v) FBS and 10mM HEPES buffer (Sigma-Aldrich). A549 cells were then rinsed with PBS and the bacterial suspension at specified OD’s were added, followed by centrifugation of 2000 x g for 5 min at RT. After each time point of incubation under the condition of 37°C with 5% CO_2_, the infected cells were fixed by 4% PFA for 30 min and washed with PBS. The cover slip was picked up from 24-well plate and mounted with DABCO containing DAPI and subjected to fluorescence microscopy (TIRF). Microscopy acquisition was performed using a Leica DMi8 microscope with a sCMOS DFC9000 (Leica) camera and a SpectraX light source (Lumencor). Phase-contrast images were acquired using transmission light (100 ms exposure) and still fluorescence images were acquired with 700 ms exposure. The Leica DMi8 filters set used were as followed: DAPI (Ex: 395/25 nm, Dc: 425 nm, Em: 460/50 nm), RFP (Ex: 545/20 nm, Dc: 562 nm, Em: 605/60 nm), and GFP (Ex: 470/40 nm Chroma ET470/40x, BS: LP 498 Leica 11536022, Em: 520/40 nm Chroma ET520/40 m). All microscopy images were processed using FIJI v.1.52q (fiji.sc).

### Infection conditions

1 ml of A549 cells in DMEM culture medium at a density of 2x10^5^ cells/ml were seeded onto each well of cell-culture treated 24 well trays (Corning Costar). The day after seeding, A549 cells in each well were rinsed with 1x PBS (pH 7.4) and 1ml of RPMI 1640 without phenol red (Gibco™) supplemented with 1% (v/v) FBS was added. The following day, *S. pneumoniae* strains were grown in C+Y medium, and upon reaching OD_600_ ∼0.2 (exponential phase), bacteria were spun down and resuspended in infection medium, RPMI 1640 without phenol red supplemented with 1% (v/v) FBS and 10mM HEPES buffer (Sigma-Aldrich). In the meantime, confluent A549 monolayers in the cell culture treated 24 well trays were rinsed twice with 1x PBS (pH 7.4). Pneumococcal suspension in infection medium was added onto the epithelial monolayer at a multiplicity of infection, MOI ∼20 (20 pneumococcal cells per epithelial cell). To optimize cell-to-cell contact, soft centrifuging was employed (400 ×g, 5 min, 4°C), subsequently, the infection culture was incubated in humidified 5% (v/v) CO_2_ atmosphere at 37°C for 2h (56).

### Cell sorting and 10x scRNA-seq

A549 cells were seeded in cell-culture treated 24 well trays (Corning Costar) as described above, and serotype 19F pneumococcal strains were added to each well at a MOI ∼20 for 2h in a humidified 5% (v/v) CO_2_ atmosphere at 37°C. Subsequently, all wells were rinsed twice with 1x PBS (pH 7.4) to remove unadhered pneumococcal cells. Cells were then detached using 200μl of 15mM of sodium citrate tribasic dihydrate (Sigma-Aldrich) per well for 5 minutes. Per condition, quadruplicate infections were performed and pooled together before detachment and passing through a 35μm strainer staining with 0.1μg/ml DAPI for 1-2 minutes on ice in the dark. Cells were then sorted using a FACSAria III Cell Sorter (BD Biosciences), equipped with a 100μm nozzle. A549 cells were gated based on FSC, SSC and DAPI, to exclude dead cells, and then sorted based on GFP expression (Fig. S1). Flow rates and dilutions were adjusted to keep the efficiency of sorting as high as possible (at a minimum above 85%). At least 30,000 GFP negative and GFP positive cells per condition were sorted into 2ml round bottom tubes (Eppendorf), that were coated prior overnight with 1% BSA in 1x PBS (pH7.4) at 4°C, containing 100μl 1x PBS (Ph 7.4) supplemented with 10% FBS. Sorted cells were enumerated using a haemocytometer (Abcam) and 1-10 dilutions of Trypan Blue solution 0.4% (Gibco), before 4°C centrifugation at 300g for 5 minutes. The supernatant was removed and cells were then resuspended in 1x PBS supplemented with 10% FBS to a concentration of 1000 cells/μl and placed on ice. Samples were then processed for and subjected to scRNA-seq using a Chromium Single Cell 3’ Reagent Kit v3 (10x Genomics), as previously described (57).

Analyses were performed in R (v. 4.4.1) with Seurat (v. 5.4.0) (58), following standard pipeline instructions and recommendations, and generally using default parameters. Briefly, cells were filtered to have a minimum of 1,500 expressed genes and mitochondrial DNA (mtDNA) contents between 0.2% and 14%, prior to regressing out the effects of those factors along with percentage of reads mapped to ribosomal, heat-shock protein and histone genes, and cell cycle stage. Data from different samples were integrated with the CCAIntegration method, and its first 15 dimensions were used for nearest neighbor graph construction. The number of identified clusters in this cell line was kept stably and conservatively low by setting the pruning parameter to 0.01 and k.param to 40, and the cluster resolution to 0.1. Differentially regulated genes were subsequently found with the FindMarkers functionality. The data from Röwekamp, Maschirow & Rabes et al. (36) were obtained from NCBI GEO accession GSE236344 and analyzed in the same fashion as described above, but using the first 20 integrated dimensions, default FindNeighbors and FindClusters parameters, and additionally filtering out cells with mtDNA > 10% and number of genes < 150 to match the analysis performed in the original study. Only WT mouse samples with *Streptococcus pneumoniae* 12 hours post infection or PBS control treatment were included. Global gene markers S100a8, Cd79a and Lamp3/Sftpb were used to respectively identify clusters of neutrophils, B cells and alveolar epithelial cells, as in the original study.

### RNA FISH

A549 and Detroit 562 cells were infected with serotype 19F strains as described above in 24 well trays (Corning Costar) at a MOI ∼20, except that in each seeded well a 12mm coverslip (Menzel^TM^) was added before seeding the epithelial cells, to allow attachment to the coverslips. After 2h, cells were washed with 600μl 1x PBS and then fixed with 3.7% formaldehyde solution for 10 min at RT. Cells were then washed 2x with 600μl 1x PBS and permeabilized in 600μl of 70% ethanol for 1-3 days at 4°C. The infected, fixed and permeabilized cells on 12mm coverslips were then hybridized to the CXCL2-, NFKBIA- and GAPDH-quasar 670 probes, following the manufacturer’s instructions available online at www.biosearchtech.com/stellaris protocols, except that instead of 1ml and 100μl volumes, 600μl and 60μl volumes were used respectively in 24 well trays (Corning Costar®). The custom Stellaris® FISH Probe set was designed against CXCL2 and NFKBIA by utilizing the Stellaris® RNA FISH Probe Designer (Biosearch Technologies, Inc., Petaluma, CA) available online at www.biosearchtech.com/stellarisdesigner. The custom Stellaris® FISH Probe set recognizing GAPDH and labelled with Quasar 670 was ordered readymade. Microscopy acquisition was performed using a Leica DMi8 microscope with a sCMOS DFC9000 (Leica) camera and a SpectraX light source (Lumencor). Z-stacked fluorescence images were acquired with 700 ms exposure. The Leica DMi8 filters set used were as followed: *DAPI (Ex: 500, Dc: 520, Em: 535), Quasar 670 (Ex: 550, Dc: 570, Em: 576)* and GFP (Ex: 470/40 nm Chroma ET470/40x, BS: LP 498 Leica 11536022, Em: 520/40 nm Chroma ET520/40 m). Images were processed using LasX v.3.4.2.18368 (Leica). All microscopy images were processed using FIJI v.1.52q (fiji.sc).

### Zebrafish husbandry and maintenance

Transparent *mitfa^w2/w2*; mpv17^b18/b18 (casper) adult zebrafish and Tg(mpx:GFP) casper transgenic fish, in which neutrophils express green fluorescent protein, were maintained at 28°C in aerated 3.6 L housing tanks under a 10 hour dark and 14 hour light cycle. Embryos were generated by natural spawning, collected shortly after fertilisation, and incubated at 28°C in E3 medium containing 5.0 mM NaCl, 0.17 mM KCl, 0.33 mM CaCl₂·2H₂O, and 0.33 mM MgCl₂·6H₂O, supplemented with 0.3 mg/L methylene blue. All zebrafish procedures were carried out in accordance with institutional and national animal welfare guidelines. Animal experimentation at UNIL complies with Swiss regulations (canton Vaud, licence VD-H28) under the Animal Welfare Act (SR 455) and Animal Welfare Ordinance (SR 455.1).

### Zebrafish infection experiments

*S. pneumoniae* 19F WT and 19F *ΔcomCDE* strains were cultured in C+Y medium to mid-logarithmic growth phase (OD595 0.2 to 0.3), collected by centrifugation at 6,000 rpm for 10 minutes, and resuspended in 0.25% (w/v) amaranth solution to facilitate visualisation during microinjection. Before injection, 2 days post-fertilisation embryos were manually dechorionated when required and anaesthetised in 0.02% (w/v) Tricaine. Embryos were randomly allocated to experimental groups. Approximately 1 nL of bacterial suspension was injected into the hindbrain ventricle of 2 dpf embryos. The number of colony-forming units (CFU) per injection (inoculum) was determined by quantitative plating of the injection volume. Inoculum doses were optimised for each experimental endpoint: a higher inoculum of approximately 2,000 CFU was used for gene expression analysis to ensure robust transcriptional responses across pooled embryos, approximately 1,000 CFU for live imaging experiments to ensure timely neutrophil recruitment over the imaging period, and approximately 300-400 CFU for survival experiments to allow mortality to develop over a meaningful time course. Across experiments, the wild-type inoculum was comparable to or slightly lower than that of mutant strains, excluding differences in starting dose as a cause of attenuation. Following injection, embryos were maintained at 28°C in 6-well plates containing egg water prepared with 60 mg/mL sea salts. For COX-2 inhibition experiments, infected embryos were treated with celecoxib (Sigma-Aldrich) prepared in DMSO or vehicle control (DMSO) at the indicated concentrations and added directly to the egg water. Survival was assessed by recording live and dead embryos at defined intervals between 24 and 72 hours post-infection. All experiments were conducted in three independent biological replicates. Survival curves were generated using GraphPad Prism 10.0 and analysed with the log-rank Mantel-Cox test, with *p* < 0.05 considered statistically significant.

### Quantitative RT-qPCR

For zebrafish gene expression analysis, total RNA was isolated from embryos collected at 4 and 8 hours post-infection using 20 embryos per biological replicate. Embryos were anaesthetised in 0.02% Tricaine prepared in egg water, transferred to 2 mL microcentrifuge tubes, and residual liquid was removed prior to lysis. Samples were homogenised in RLT lysis buffer from the RNeasy Mini Kit (Qiagen; 74104) by repeated passage through a 27-gauge needle attached to a 1 mL syringe until complete disruption was achieved. Total RNA was purified using the RNeasy Mini Kit (Qiagen) according to the manufacturer’s instructions, including on-column DNase digestion to remove genomic DNA. Complementary DNA was synthesised from purified RNA using the High-Capacity cDNA Reverse Transcription Kit with RNase inhibitor (Thermo Fisher Scientific; 4374966) following the supplier’s protocol. Primers were designed to span exon-exon junctions using NCBI Primer-BLAST and are listed in Supplementary table 2. Quantitative PCR reactions were prepared using PowerSYBR Green PCR Master Mix (Thermo Fisher Scientific; 4367659), 0.5 µM of each primer, and 1 µL of cDNA template in a final volume of 10 µL. Reactions were performed in technical triplicate on a QuantStudio 5 Real-Time PCR System using standard cycling parameters. Gene expression levels were normalised to the reference gene *bactin1* (β-actin 1), and relative expression was calculated using the ΔΔ*Ct* method (59).

### Time-lapse fluorescence imaging of zebrafish embryos

Time-lapse imaging of *Tg(mpx:*GFP) zebrafish embryos was performed using a Nikon AXR confocal microscope. Embryos were infected at 3 days post-fertilisation by injection of 1,000 CFU of *S. pneumoniae* 19F WT or 19F Δ*comCDE* into the hindbrain ventricle. Imaging was initiated at 1.5 hours post-infection and images were acquired at 20-minute intervals for 3 hours. Following injection, anaesthetised embryos were embedded in 0.5% low-melting point agarose dissolved in egg water containing 60 mg/mL sea salts and supplemented with 0.02% (w/v) Tricaine. Embryos were mounted in open uncoated 8-well chamber slides and maintained at 28°C in a temperature-controlled imaging chamber. NIS Elements and ImageJ software were used for image processing and neutrophil quantification.

## Supplementary information

Supplementary Information is available for this paper. Supplementary figures and tables are provided in a separate Word document.

## Data availability

Processed data are provided as a Supplementary Data Excel file. Single-cell RNA sequencing data generated in this study have been deposited in the European Nucleotide Archive (ENA) under accession number E-MTAB-16871. RNA fluorescence in situ hybridization images have been deposited in the BioImage Archive under accession number S-BIAD3076. The mouse lung scRNA-seq data reanalysed in this study are publicly available from the NCBI Gene Expression Omnibus under accession number GSE236344.

## Author contributions

V.M. conceived the study, designed and performed experiments including scRNA-seq, smFISH, luciferase and fluorescence microscopy assays, analysed data, and wrote the original manuscript draft. V.d.B. performed the scRNA-seq analyses and contributed to manuscript editing. K.K.J. designed and performed zebrafish infection, time-lapse imaging, and survival experiments, and contributed to manuscript editing. J.K. conceived the study and designed epithelial fluorescent lines. J.P. performed bioinformatics analysis of scRNA-seq data and contributed to data interpretation. M.R.G. designed the quadGreen fluorescent cassette. C.C.M. contributed to scRNA-seq methodology and data interpretation. R.A. provided conceptual input and contributed to manuscript editing. B.D. provided bioinformatics infrastructure and supervision for scRNA-seq analyses and contributed to manuscript editing. J.W.V. conceived and supervised the study, acquired funding, and contributed to manuscript writing and editing. All authors reviewed and approved the final manuscript.

## Acknowledgements

The authors would like to thank the Genomics Technology Facility (GTF) at the University of Lausanne (UNIL) for their support with single-cell RNA sequencing library preparation and sequencing. We also thank Sven Hammerschmidt for kindly providing strain EF3030. V.d.B was supported through a SNSF PostDoc Mobility fellowship (P500PB_225439). Work in the lab of J.W.V. was supported by the EU project NOSEVAC and by the SNSF grants 320030-236203, 320030-231669 and NCCR 51NF40_180541. The funders had no role in study design, data collection and analysis, decision to publish or preparation of the manuscript. Portions of the introduction, results, and discussion were edited for clarity and language with the assistance of Claude (Anthropic), an AI language model. All scientific content, interpretations, and conclusions are the authors’ own.

