## Supplementary Information for "Epithelial innate immune sensing of pneumococci is inherently restricted to a small cellular minority across species and infection niches"

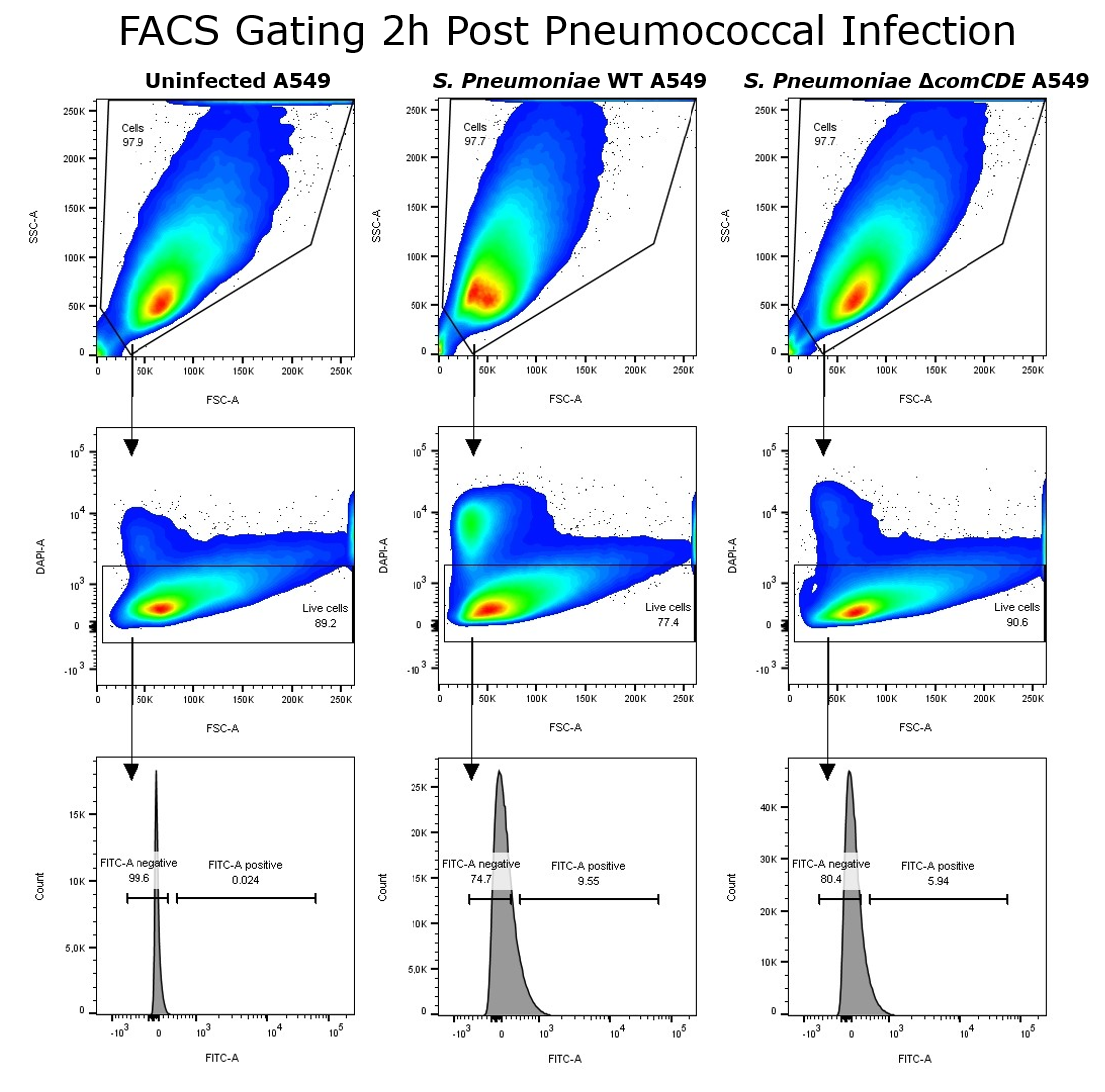

Supplementary Fig 1. Gating strategy used in the FACS Aria III to sort out single A549 lung epithelial cells that had attached *S. pneumoniae* (see materials and methods for more details).

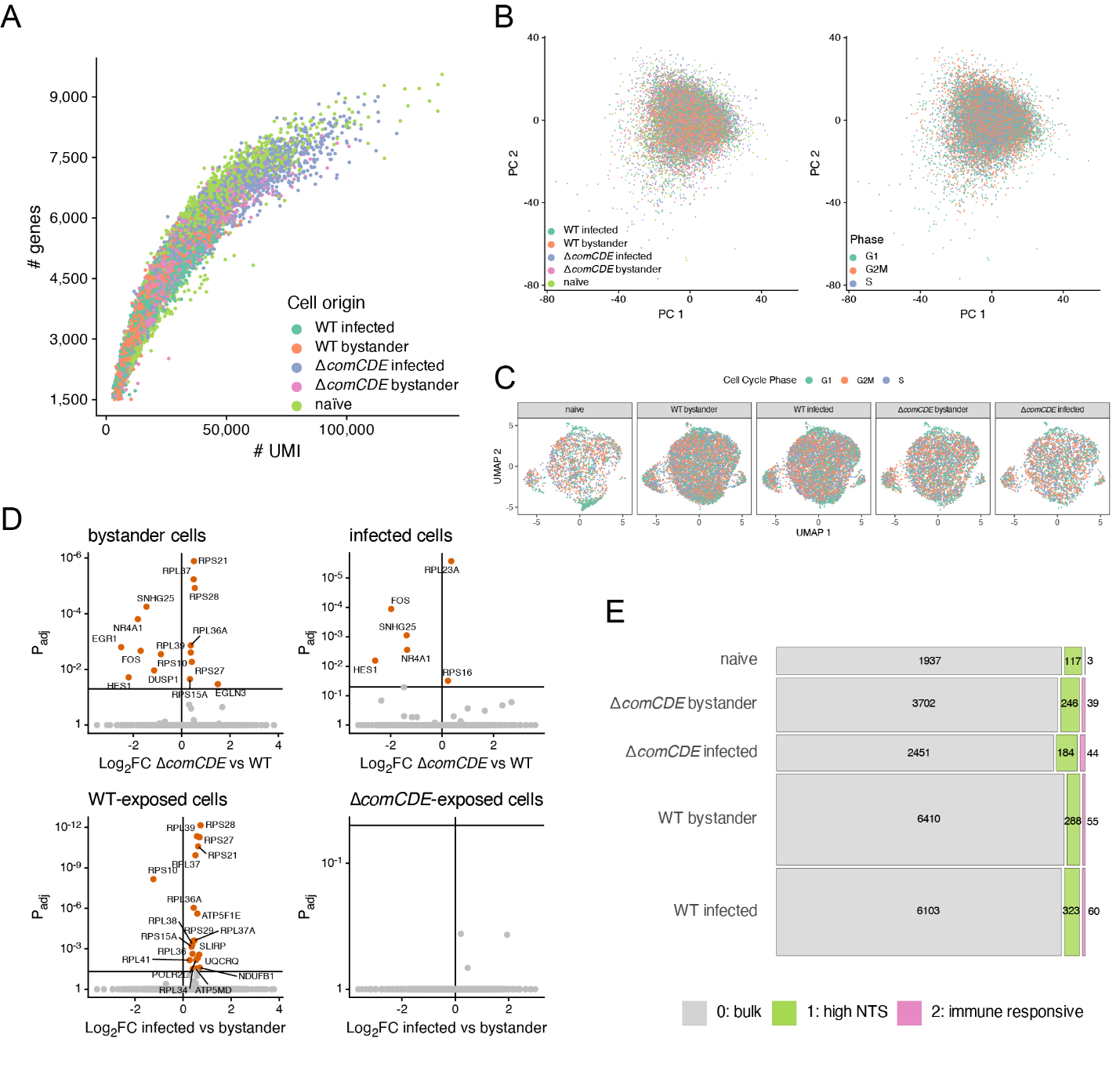

Supplementary Fig 2. scRNA-seq data exploration. A) Numbers of unique molecular identifiers (UMI) and genes for which transcripts were detected per cell. B) Principal Component Analysis of integrated transcriptomes colored by either sample of origin or scored cell cycle phase. C) Uniform Manifold Approximation and Projection of integrated transcriptomes split by sample of origin. D) Differential expression analyses of pair-wise sample contrasts, as indicated per volcano plot. Cutoffs: 0.05 for FDR-adjusted p-values and 0 for log2 fold change. E) Mosaic plot displaying numbers of retained cells used in downstream analyses per combination of sample of origin and identified cluster

Supplementary Table 1. List of bacterial strains and plasmids used in this study

| **Name** | **Genotype** | **Reference** |
| --- | --- | --- |
| VL2539 | *S. pneumoniae* D39V, *bgaA::PssbB*_quadGreen | This study |
| VL2540 | *S. pneumoniae* D39V, *hlpA-*mCherry, bgaA*::PssbB_quadGreen* | This study |
| VL3889 | *S. pneumoniae* 19F, *ssbb*::*ssbb*-*luc*-*kan* | This study |
| VL3894 | *S. pneumoniae* 19F, *ssbb*::*ssbb*-*luc*-*kan*, *comCDE*::*ery* | This study |
| VL3964 | *S. pneumoniae* 19F, *bgaA*::*Plac*-quadGreen-*spec* | This study |
| VL3990 | *S. pneumoniae* 19F, *bgaA*::*Plac*-quadGreen-*spec, comCDE*::*ery* | This study |
| VL3668 | *S. aureus* NCTC 8325 | Lab collection |
| VL995 | *E. coli* Stbl3 | Lab collection |
| pVL306 | pJWV102 | Lab collection |
| pVL2129 | pPEPZ-Plac-msf-GFP-DSM, spc^R^ | Lab collection |
| pVL2130 | pPEPZ-Plac-msf-GFP-OPT, spc^R^ | Lab collection |
| pVL2132 | pPEPZ-Plac-mNeonGreen-Opt, spc^R^ | Lab collection |

Supplementary Table 2. List of primers used in this study

| **Name** | **Sequence** |
| --- | --- |
| OVL96 | cttgccacgaaaaaagttgc |
| OVL167 | TGGTGATGACACCGTCTTTG |
| OVL1582 | CACGTTCACCTGCGAGAGACAAACATGTCAAAAGGAGAAGAA |
| OVL1583 | CACGTTCACCTGCGAGACTTAATTATTATTTATAAAGTTCGTCCATACCG |
| OVL1584 | CACGTTCACCTGCAGCAAACTCGAGAAAAAAAAACCGCGCCCCT |
| OVL1585 | CACGTTCACCTGCAGCATGTCCTCCTTTATTATTTGTATAGTTCGTCCATGC |
| OVL1586 | CACGTTCACCTGCCTACAGGAGGCAAATATGTCAAAAGGAGA |
| OVL1587 | CACGTTCACCTGCCTACAGTTATTACTTATAAAGCTCATCCATGC |
| OVL1588 | CACGTTCACCTGCCCTTTAAGGAGGCAAATATGTCAAAAGGC |
| OVL1589 | CACGTTCACCTGCCCTTTCCTTAATTATTATTTGTATAGTTCGTCCATGC |
| OVL2870 | TGTTTGAGCCTGAAACTAG |
| OVL5539 | ACGATACCTTGAGTGACAG |
| OVL5643 | gcagtcacagcaagtgttag |
| OVL5644 | accgtctgcacttcactaag |
| OVL5653 | TGACGTCACCTGCCTTGGCtcgataaggacaatcaaa |
| OVL5654 | TGACGTCACCTGCATGGgaGCAAAAAACTGGACGGAGTCCCGT |
| OVL5655 | TGACGTCACCTGCATGGccAAAAAAAAACCGCGCCCTGTCAGG |
| OVL5656 | TGACGTCACCTGCCCTATTggaggttattggccataa |
| OVL5657 | gactttatgcaccgtaagtg |
| OVL5658 | tgaacaggctatcaatgag |
| OVL10524 (nfkbiz_qPCR_FW) | TGACGGATACACACCGATGGA |
| OVL10525 (nfkbiz_qPCR_REV) | TCCTGCTGGATCTGCCATTG |
| OVL10526 (tnfaip3_qPCR_FW) | CTCAGAACCAACGGAGATGGG |
| OVL10527 (tnfaip3_qPCR_REV) | AACCCATTCCTCCTCCCAGT |
| OVL10528 (ccl20b_qPCR_FW) | GAGACCGAGTCCGCGATATG |
| OVL10529 (ccl20b_qPCR_REV) | TAAGACCCGTTCTTGCGTCC |
| OVL10530 (cxcl8a_qPCR_FW) | GAGCTTGAGAGGTCTGGCTG |
| OVL10531 (cxcl8a_qPCR_REV) | TTGTCATCAAGGTGGCAATGATCT |
| OVL10532 (efna1a_qPCR_FW) | GAGTTTCGACAGGGCGAGAG |
| OVL10533 (efna1b_qPCR_REV) | GAGCCACAACAGGATCATCTGC |
| OVL10534 (efna1b_qPCR_FW) | TCCCAGGAGGCAGAACGATA |
| OVL10535 (efna1b_qPCR_REV) | TGGTGGAGAGGCTTGGAGAT |
| OVL10536 (nfkbiaa_qPCR_FW) | CCTAACTACAGCGGACACACG |
| OVL10537 (nfkbiaa_qPCR_REV) | CAGGTTCTGCAGGTCTACGG |
| OVL10538 (nfkbiab_qPCR_FW) | AGTCTGAGGTGGAAGACCTGT |
| OVL10539 (nfkbiab_qPCR_REV) | GTTTTCAGCCGCCTGATGTG |
| OVL10540 (ptgs2a_qPCR_FW) | GTCCTATTACACCCGCACCC |
| OVL10541 (ptgs2a_qPCR_REV) | TGCCCCAGATCCACTCCAT |
| OVL10542 (ptgs2b_qPCR_FW) | CGGCATAATGCGATACGTGC |
| OVL10543 (ptgs2b_qPCR_REV) | TGGCAGCTCTTTCTTACCTGC |
